# Defective Hippocampus-Dependent Spatial Memory in Mouse Model of Polyendocrine metabolic ovarian syndrome (PMOS, formerly PCOS)

**DOI:** 10.64898/2026.04.30.721991

**Authors:** Samanwitha Rao, Betcy Susan Johnson, Malini Laloraya

## Abstract

Polyendocrine metabolic ovarian syndrome (PMOS), formerly called as Polycystic Ovarian Syndrome (PCOS), is a complex endocrine disorder characterised by hyperandrogenism, oligo-or anovulation, and polycystic ovaries. Endocrine dysfunction in PMOS disrupts both hormonal and neurotransmitter balance, contributing to the psychological distress frequently reported by affected individuals. Although hormonal imbalances have been associated with memory impairments, their specific contribution to cognitive dysfunction in PMOS remains incompletely understood. In this study, we investigated the impact of PMOS on the hippocampus, a brain region critical for memory formation and highly sensitive to sex steroid modulation. A dehydroepiandrosterone (DHEA)-induced PMOS mouse model was employed to assess anxiety-like behaviour, locomotion, and memory. In the open field test (OFT), DHEA-treated mice spent significantly less time in the central zones and travelled a shorter total distance compared with controls, indicating increased anxiety-like behaviour. DHEA treatment also resulted in significantly impaired performance in both the object location test (OLT) and novel object recognition test (NORT), as reflected by a reduced discrimination index. Analysis of hippocampal immediate early gene expression using qRT-PCR revealed altered transcription of memory-related markers, including downregulation of *Npas4* and *Grin2a*, and upregulation of *Grin1*, *Arc*, *Egr1*, and *Egr2*. Collectively, these findings suggest that elevated androgen levels induce anxiety- and depression-like behaviours and impair cognitive function, including spatial, recognition, and motor learning abilities, in PMOS. Our results further indicate that disrupted cortex-hippocampus communication may underlie these cognitive deficits, underscoring the importance of evaluating memory and cognitive health in women with PMOS to support brain health and overall well-being.

**Graphical summary:** Cartoon representation of the impact of hyperandrogenism on anxiety, spatial memory, and cognition in PMOS pathology.

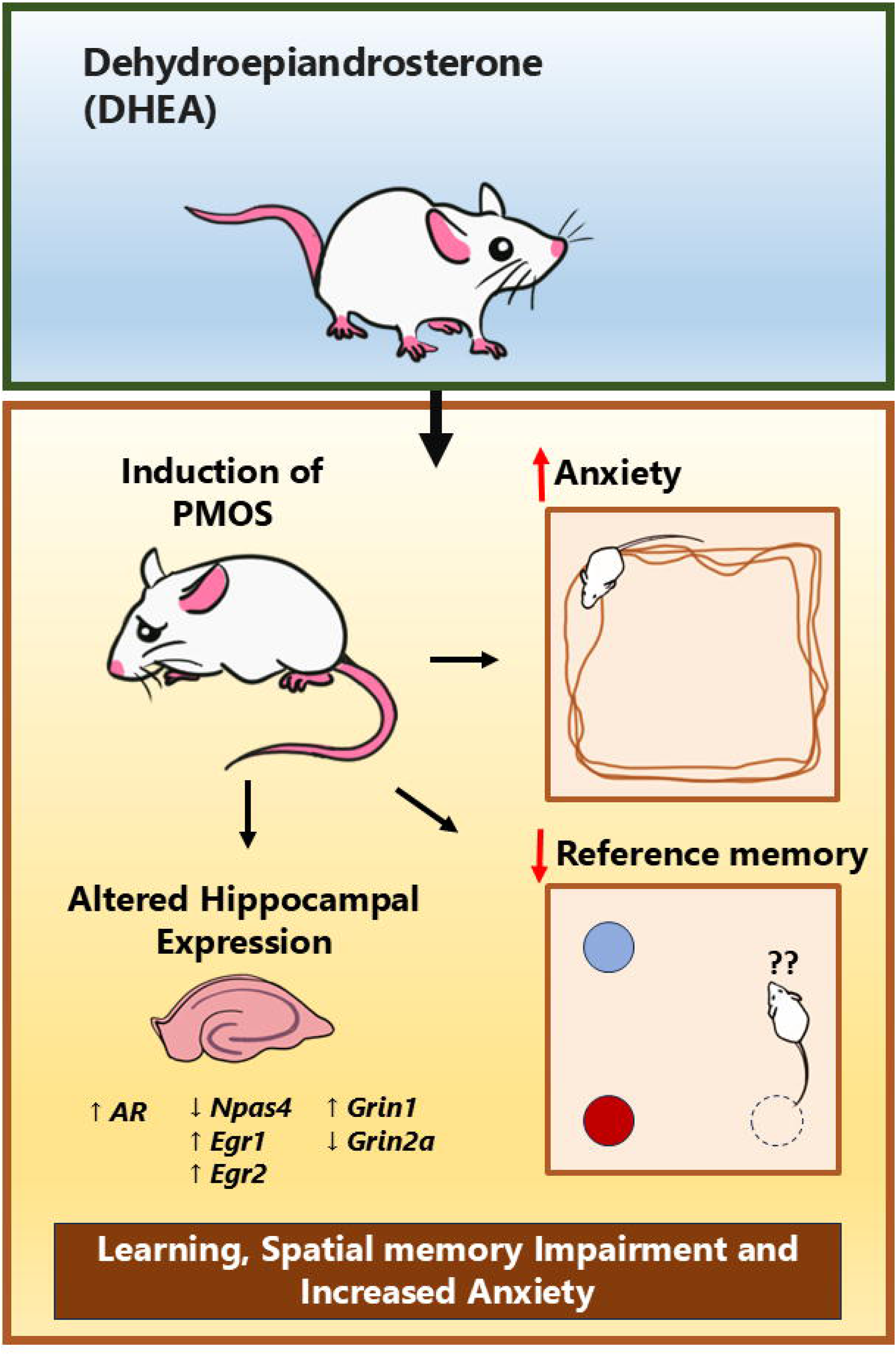

## Introduction

Polyendocrine metabolic ovarian syndrome (PMOS) (formerly Polycystic ovarian syndrome PCOS), is a multifactorial disorder linked to female infertility(1). PMOS was first defined in 1935 as a case of cystic ovaries with amenorrhea presumed to be caused by abnormal endocrine function(2). Subsequently, clinical studies have established that elevated androgen levels are a prominent feature in individuals with PMOS. As per Rotterdam criteria, diagnosis of PMOS can be ruled when a patient presents two of these three symptoms: hyperandrogenism, oligo-anovulation, and polycystic ovaries(3). By this criterion, the global prevalence of PMOS worldwide is 5-20% with 4.8% among the Caucasian population in the United States, in 8-12% Caucasian population in the rest of the world(4) and a much higher prevalence of 19.6% in Indian women. The additional effects of PMOS on the metabolism gained traction after the high incidence rate of hyperinsulinism and insulin resistance in hyperandrogenic women was discovered(5). PMOS has also been linked with dyslipidemia, cardiovascular risk, obesity, and type 2 diabetes mellitus(6).

Beyond metabolic effects, PMOS significantly impacts the psychological well-being by increasing distress and therby reducing quality of life(7–9). PMOS has been found to be linked to psychiatric syndromes due to observed depressive episodes and social phobia as well as higher suicide attempts(10). Atleast one psychiatric disorder including mood disorder , major depression and bipolar disorder has been reported in 57% of PMOS women ((11). A large Swedish cohort, reaffirmed psychiatric disorders in PMOS women with an increased risk to bipolar disorder, depression, anxiety, schizophrenia, with highest odds ratio for autism spectrum disorder and tics(12). An fMRI (functional magnetic resonance imaging) study shows that PMOS patients have altered brain activity in the middle frontal gyrus, associated with depression. Women with PMOS are reported to exhibit altered white matter microstructures, characterized by a higher volume fraction in the corpus callosum and decreased axial diffusivity in white matter fibers(13) alongwith lowered cognitive performance than controls, independent of education and age. In a recent larger CARDIA cohort, PMOS women exhibited reduced cognitive and attention performance on Stroop test, Rey Auditory Verbal Learning Test (RAVLT) (verbal learning and memory), and category and letter fluency tests (semantics and attention) as well as lower white matter integrity(14).

The impaired hypothalamic-gonadal axis is believed to contribute to the abnormal endocrine function in PMOS(15). A hyperactive GnRH pulse secretion from the brain leads to abnormal pulsatile luteinising hormone (LH) secretion(16). Interestingly, high LH levels are associated with a poor ability to recall memories(17). In PMOS women, high plasma LH levels are associated with reduced right frontal lobe functional connectivity, potentiall interfering with cognitive functions, including visuospatial working memory, face processing, and episodic memory in women with PMOS(18).

Sex steroid hormones, such as testosterone and oestrogen, also exert significant effects on cognition(19). Women are more proficient in verbal skills than men, who tend to perform better in spatial ability tests(20). This spatial ability performance advantages observed between sexes have been linked to higher levels of testosterone in males although testosterone supplementation does not improve cognition in older men or women(21). Cross-sectional observational, epidemiological, and retrospective studies have found that PMOS women report less efficient executive functioning and utilized more neural resources during a working memory task than those without PMOS and thus a greater risk of cognitive decline (22). Another study reported that higher levels of free testosterone in PMOS might be associated with lower outcomes on neuropsychological tasks like verbal and visuospatial memory as well as verbal fluency and manual deftness(23). Women with PMOS scored higher on three-dimensional mental rotation tasks which were positively associated with circulating testosterone(24). Although mental rotation is traditionally associated with the parietal lobe, research suggests that it is modulated by the hippocampus in a sex-dependent manner(25). This is based on reports that males have a larger anterior hippocampus and gray matter volume of this area correlates significantly with 3D mental rotation score(25). The hippocampus plays a central role in memory formation and spatial navigation(26; 27).

The cognitive and neural effects of testosterone suggest a significant role for androgen signalling within the brain Hippocampus functions are influenced by sex hormones, and this is substantiated by the presence of androgen receptors in extranuclear sites of all hippocampal sub-regions including CA1 pyramidal cell nuclei and punctate processes making it a key area of interest in PMOS(28). Remarkably, the loss of neuron-specific androgen receptors (ARs) protects from PMOS, even in hyperandrogenic conditions(29) which underscores the importance of neuronal AR in PMOS pathogenesis. In the Dehydroepiandrosterone (DHEA)-induced PMOS mouse model, signs of anxiety and depression-like behaviour have previously been demonstrated by reduced movement in the open field test (OFT) and increased immobility during the forced swim and tail suspension tests(30). Yu et al further linked depression-like behavior due to reduced brain monoamines and/or their metabolites implicating the contribution of hyperandrogenism to the psychological symptoms of women with PMOS.

To this date, no study has directly examined PMOS-induced hippocampal-dependent memory impairments in DHEA-treated mice models nor tested whether these deficits co-occur with depression-and anxiety-like behaviours. In this study, we investigated whether DHEA-treated mice model of PMOS show hippocampal-dependent memory deficits. To this end, we employed the Object Location Test (OLT) and Novel Object Recognition Test (NORT), which assess reference memory through an animal’s recognition of spatial or object-based changes in its environment(31). The OLT probes spatial memory dependent on the hippocampus, whereas the NORT evaluates recognition memory, which relies on interactions between the hippocampus and adjacent cortical areas(31). Both tasks minimise stress, thereby reducing stress-related confounding effects in the DHEA model(32). The behavioural tests were complemented with molecular analysis of early-response and long-term memory markers in hippocampus. An overview of the methodology is provided in Figure 1.

**Figure 1:**
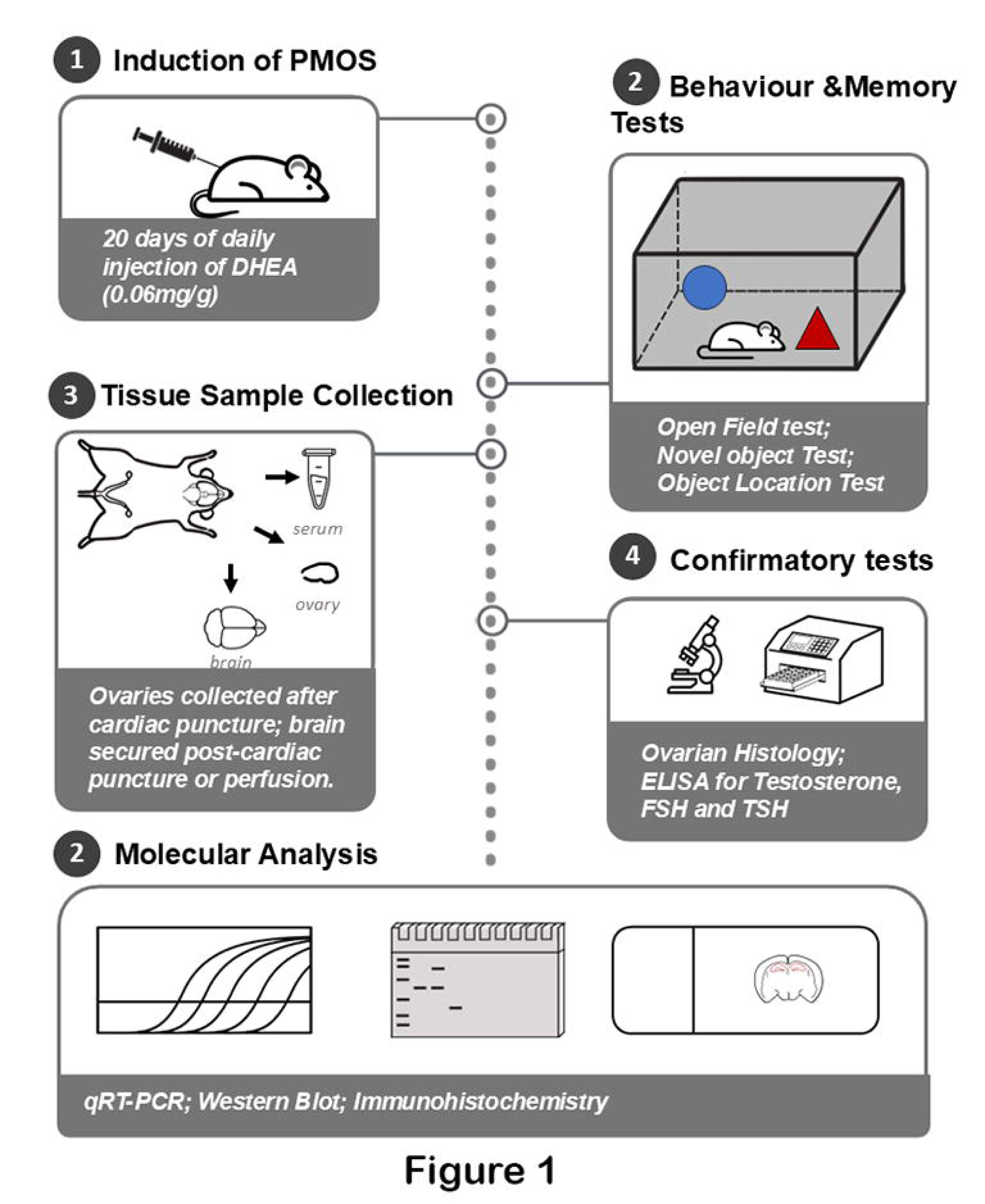
Schematic representation of the experimental design, including: A) induction of PMOS using daily subcutaneous DHEA injections for 20 days; B) behavioural and memory assessments comprising the OFT, NORT and OLT; C) tissue sample collection, including serum, ovary and brain extraction following cardiac puncture or perfusion; D) confirmatory analyses involving ovarian histology and serum hormone measurements (testosterone, LH and FSH); and E) molecular analysis workflow depicting qRT-PCR, Western blotting and immunohistochemistry performed on collected tissues.

## Materials and Methods

### Animal handling and surgery

This study received approval from the Rajiv Gandhi Centre for Biotechnology, Animal Ethical Committee (IAEC/797/MAL/2020). Animals for these experiments were housed at the Animal Research Facility of RGCB and received food and water *ad libitum* in controlled temperature conditions with a 14:10 light-dark cycle. Swiss albino mice were chosen for this study based on previous work in our laboratory validating their suitability for DHEA-induced PMOS models as it mimics the polycystic ovaries and abnormal serum hormone levels observed in PMOS patients(33). Also swiss albino mice have been previously used for assessing spatial memory using OLT(34) and object recognition task(NORT) with no difference in strains based on discrimination index(35; 36). Prepubertal (25-30 days old) Swiss Albino female mice were treated with DHEA, and animals that exhibited elevated androgen levels following treatment were included in the experimental group (n = 15), while untreated mice were maintained as controls (n = 22). For 20 consecutive days, the DHEA-treated group received a subcutaneous injection of DHEA (6mg/ 100g body weight) dissolved in 0.1 ml propylene glycol. The control group was instead administered 0.1 ml propylene glycol. Mice were weighed on the initial day of the experiment and on the day they were euthanized.

### Vaginal Cytology

Daily vaginal smears were collected using PBS lavage and examined under a brightfield microscope to determine the oestrous stage. Prolonged oestrus confirmed successful PMOS induction (Figure 2G).

**Figure 2:**
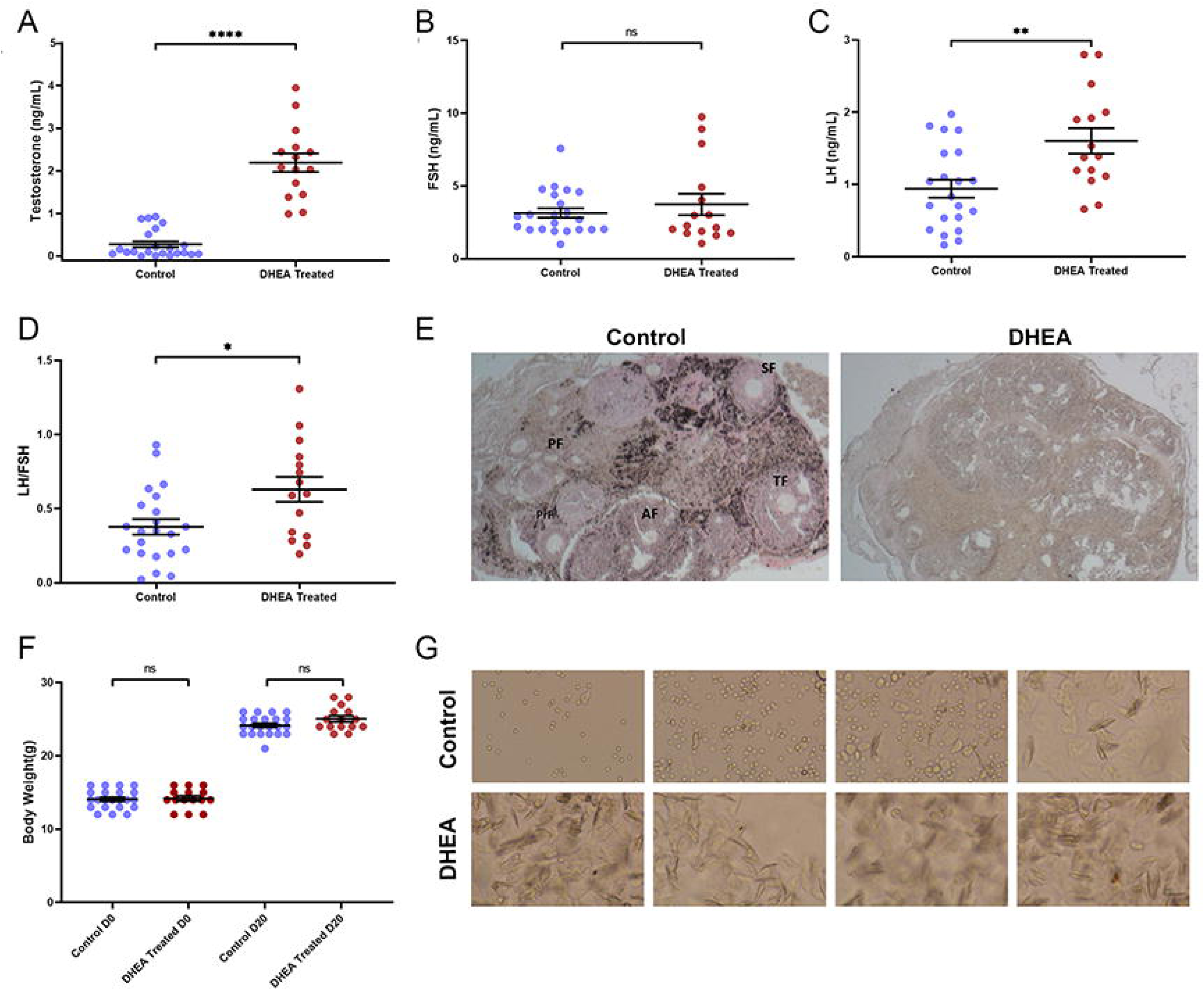
Assessment of PMOS-like traits in DHEA-treated PMOS mice: Serum levels of A) testosterone, B) FSH, C) LH, D) LH/FSH ratio in control(n=22) and DHEA-induced PMOS (n =15) mice, E) Depictive image of ovary sections of control and DHEA induced PMOS mice stained using Hematoxylin and Eosin marking the Primordial follicle (PrF), primary follicle (PF), secondary follicle (SF), antral follicle (AF), and tertiary follicle (TF), F) Graphical representation of body weight of control and DHEA induced PMOS mice at the experiment initiation day and the end of the experiment, G) Depictive image of vaginal smear of control and DHEA induced PMOS mice in showing stages of estrous cycle. Data presented as mean ± SEM. Control group (blue) and DHEA-treated group (brown) are shown as individual data points in the figure. The Unpaired Student’s T-test or Mann-Whitney U Test was calculated for analyzing statistical significance. P-values <0.05 were set to be significant. *P-value <0.05, **P-value < 0.01, ***P-value <0.001 and ****P-value <0.0001.

### Behavioral experiments

Behavioral studies of the mice were conducted after 20 days of treatments. All the animals were handled briefly daily till the testing day. The tests were carried out in the order of OFT (Open Field Test), NORT (Novel object recognition task) and OLT (Object Location Task) (Figure 3A). The protocol is based on the NORT-OLT protocol by Denninger et. al(37) . An acclimation period of a minimum of 1 hour in the testing room preceded every experiment. The experiments were conducted with a grey open field box (70 cm × 70 cm × 40 cm). To reduce the stress from bright lighting, all the behavioural assays were performed in a diffused lighting room with 15 - 20 lux illumination(35). All trials were videotaped to score the behavior and movement of the test subjects.

**Figure 3:**
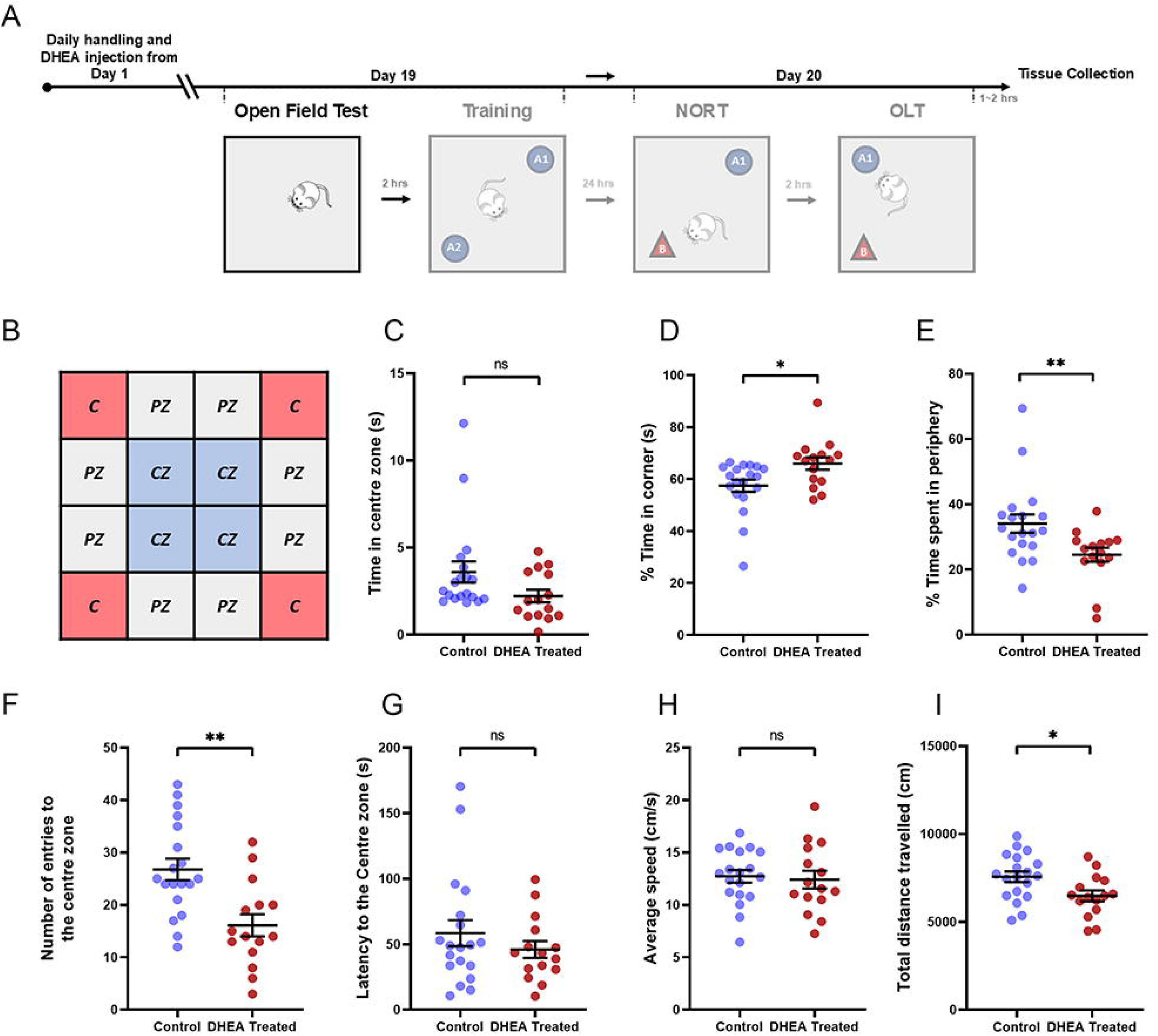
Assessment of anxiety-like behaviour and impaired cognition in DHEA-induced PMOS mice. Schematic representation of A) the behaviour protocol and B) A) Overlay of central zones (CZ), peripheral zones (PZ) and corners used to infer tracking of mice in the OFT. Performance of control and DHEA treated mice in OFT: Dot plot representations of C) time spent in central zones, D) time spent in corners, E) time spent in the periphery, F) the number of entries to the center zone, G) latency to the center, H) average speed I) total distance travelled. Control group(blue, n=19) and DHEA-treated group(brown, n=15) are shown as individual data points in the figure. Outliers have been removed. Data presented as mean ±SEM. Unpaired Student’s T-test or Mann-Whitney U was calculated for analysing statistical significance. P-values <0.05 were set to be significant. *P-value <0.05, **P-value < 0.01, ***P-value <0.001 and ****P-value <0.0001.

### Open field test (OFT)

OFT is a baseline behavioral assessment that assesses the exploratory behavior and locomotor activity of mice. The difference in time spent at the center and periphery of arenas and the total distance traveled by mice is a measure of anxiety and depression-like behavior. We assessed the relative anxiety and depression levels and locomotor activity of the DHEA-induced PMOS mice and control mice using an OFT(Figure 3A). For OFT, mice were individually introduced in randomized corners of the box and given 10 minutes to explore the arena. All trials were recorded on camera and were calculated post-experiment. The box was split into a 16-square grid using the tracking software, AnimalTA as illustrated in Figure 3B. Square grids are further divided into central zone(CZ), periphery zone(PZ) and corners(C). The CZ comprises the four central square boxes while corners are of the four square boxes at each corner as per the method detailed elsewhere(38) . The remaining square boxes on the side are considered as the PZ. The average speed, average speed moving, traveled distance, the latency to enter the CZ, no. of entries into the CZ, and time spent in three regions—the CZ, PZ, and the corners were calculated using the same. The chamber was thoroughly cleaned with 70% ethanol and dried between two tests.

### Novel object recognition task (NORT)

The NORT was also divided into three phases: habituation, training, and testing. In the Open Field Test (OFT), mice undergo habituation as they explore the empty box, acclimating to the environment before further training. In the training phase, after twenty-four hours of habituation, we positioned the mouse in the centre of the box with two objects, A1 & A2, placed in two random opposing corners and gave them 10 mins to explore the arena. After an hour, during the trial phase, the mice explored the open arena for 10 minutes in the presence of both the familiar object A1 and a novel object B. The novel object was randomised across trials to minimise object-specific bias. All objects used for NORT and OLT were made of non-toxic and odourless plastic. The criterion for exploration is active sniffing of the things within 2 cm of the object. The video recordings were reviewed, and the time spent exploring on familiar and novel objects was recorded. Trials were scored blindly. Only trials with at least 20 seconds of object exploration were included to reduce the effect of stress and anxiety-like behaviour on exploration. A schematic illustration of NORT is shown in Figure 3A. The discrimination index was calculated as:

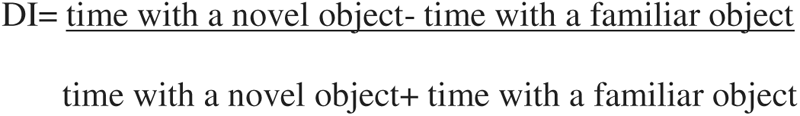

### Object location task (OLT)

The OLT was conducted after the NORT as suggested by Denninger et al. for studies that examine the hippocampus. In the object location task trial, the mice were returned to the apparatus, with the familiar object shifted to a novel spatial location, and permitted to explore for 10 minutes. The other object remains where it was during the NORT (Figure 4A). We analyzed the trial phase afterward through video recording and noted the time spent exploring the moved and unmoved object. In both tasks, objects and the chamber were cleaned thoroughly with 70% ethanol and dried between two individual tests. The trials were scored blindly. Only trials with at least 20 seconds of object exploration were included to reduce the effect of stress and anxiety-like behaviour on exploration. To determine the discrimination index, the following formula was employed:

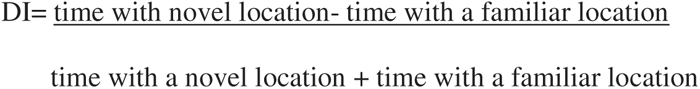

**Figure 4:**
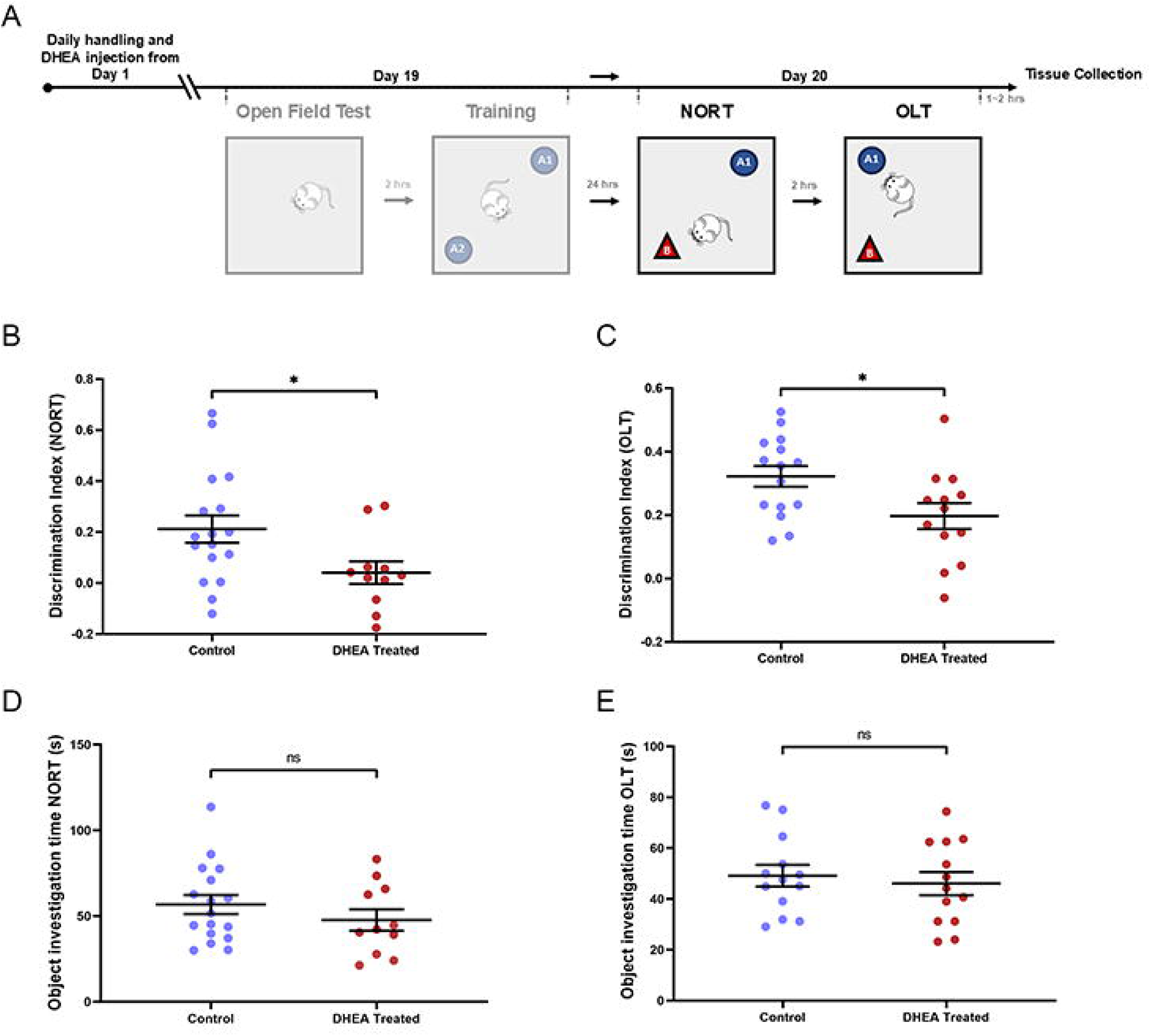
Assessment of reference memory in DHEA-treated PMOS mice vs controls. Schematic representation of A) the behaviour protocol. Dot plot depiction of the performance of control and DHEA-treated mice in B) NORT and C) OLT. Relative preference for the novel object or location was calculated by a discrimination index (DI). Fig. 4D & E represent the total time spent investigating the objects in NORT and OLT respectively. Control group (blue) and DHEA-treated group (brown) are shown as individual data points in the figure. Data presented as mean ±SEM. Unpaired Student’s T-test or Mann-Whitney U was calculated for analysing statistical significance. P-values <0.05 were set to be significant. *P-value <0.05, **P-value < 0.01, ***P-value <0.001 and ****P-value <0.0001.

### Tissue collection

After OLT, mice were subjected to CO2 euthanasia. Brain structures were removed, and the separated hippocampus for the expression studies was immediately submerged in ice-cold RNA later (Cat. #R0901,Sigma-Aldrich, USA) before being stored at −80°C. Ovaries were collected for histological confirmation of polycystic ovaries using hematoxylin & eosin stain. Blood samples were collected via cardiac puncture at the time of sacrifice to assess circulating hormone levels.

### Isolation of RNA and cDNA Synthesis

Total RNA was isolated from the bilateral hippocampi of both DHEA-induced and control group mice (n=4) using the miRNAeasy Mini Kit (Cat. # 217004, Qiagen, Netherlands) by the manufacturer’s instruction. The quantity of RNA was determined using the QuantiFluor® RNA System (Cat. #E3310, Promega, USA). cDNA was transcribed from 1µg of the extracted total RNA using SuperScript VILO™ cDNA Synthesis Kit (Cat. # 11754-050, Thermo Fisher Scientific, USA) as per manufacturer’s instructions. The amount of cDNA in the sample was quantified using QuantiFluor® dsDNA System (Cat. # E3190, Promega, USA) and diluted for subsequent real-time time-PCR.

### Quantitative Real-Time PCR (qRT PCR)

qRT-PCR primers were designed using NCBI PRIMER BLAST (Sigma-Aldrich, USA) (Supplemental Table 1). For qRT-PCR, 150ng cDNA, 0.5 µl primer pair mix and 2.5ul Power SYBR Green PCR Master Mix (Life Technologies). During the reaction cycle, the temperature was set to 95°C for 15 seconds and then to 60°C for 1 minute. Ct value was determined by monitoring changing fluorescence levels using an ABI Biosystems HT 5200. Relative gene expression was determined using the ΔCt method, normalising all target transcripts to 18S rRNA.

#### Western Blot

The hippocampus lysate was prepared using RIPA buffer (supplemented with phosphatase inhibitor and protease inhibitor cocktails (Cat. #P5726-1ML, #P00444-1ML, # S8820-20TAB respectively; Sigma-Aldrich, USA). The extracted protein was quantified using the BioRad DC Protein Assay Kit(Cat # 5000116 ; Bio-Rad Laboratories, CA, USA). Equivalent amounts of protein lysate were taken from both the PMOS and Control groups (n=3) and separated using a 10% SDS-PAGE gel. The separated proteins were subsequently transferred onto a PVDF membrane (Cat #1620177; Bio-Rad Laboratories, CA, USA). The membrane was then incubated overnight at 4°C with primary antibodies specific to NPAS4 (Cat #MA5-27592, Thermo Fisher Scientific, USA), AR (sc-7305; Santa Cruz Biotechnology, USA)), and EGR1 (Cat. #MAA416Hu21; Cloude-clone, USA). The membranes were then incubated with HRP Conjugate secondary antibodies for an hour at room temperature, followed by chemiluminescence detection and visualized using Chemidock. The band density was quantified using Image Lab software (Biorad) using Histone 3(cat. No. No. H-0164; Sigma-Aldrich, USA) bands or β actin (cat. No. # A2228; Sigma-Aldrich, USA) bands as normalization controls.

### Immunohistochemistry and Immunofluorescence

#### Tissue preparation

For immunohistochemical analysis, animals were anaesthetized using isoflurane inhalation and perfused with 1x PBS and then with 4% paraformaldehyde transcardially. Dissected brain tissues were fixed in 4% PFA overnight and cryoprotected by incubating at 4°C overnight in 30% sucrose solution with 0.1% Sodium azide. Tissues were then embedded in optimal cutting temperature (OCT) compound (Cat. #23730571; Fischer Scientific, USA), flash frozen and kept at -80°C until cryosectioning. 20µm thick coronal sections of the brain tissue containing the hippocampus area were obtained using a cryostat (CM1950, Leica Microsystems, Germany). The sections were affixed on Superfrost Plus microscope slides and preserved at -80°C.

Hippocampus sections were subsequently treated with 0.1% Triton X-100 in PBS for 1 hour, followed by exposure to 1% sodium borohydride for 20 minutes following a PBS wash lasting 10 minutes.Sections were then incubated with Rodent Block M (Cat. #RBM961; Biocare Medical, USA) for 20 min to prevent nonspecific endogenous mouse IgG binding.. After blocking, sections were incubated with NPAS4 (1:100 dilution) (Cat #. MA5-27592; Thermo Fisher Scientific, USA) overnight at 4°C. Sections were then washed three times using PBS for 10 min each and conjugated with the secondary antibody, DyLight 488 (1:500) (Cat #. 53024; Thermo Fisher Scientific, USA), by incubating for 1hr at room temperature. The sections were washed with PBS for 10 mins and were counterstained with PI (1:10000). Prolong Diamond Antifade (Cat #. P36961; Thermo Fisher Scientific, USA) was used to mount the section. The mounted sections were imaged with a confocal microscope (Olympus FV 3000, Japan). The fluorescence intensity of NPAS4 in the hippocampal Region of Interest (ROI) was quantified using Image J (39).

#### Ovarian Histology for Polycystic Ovarian Morphology

Ovaries from both the Control and DHEA treated mice were immediately paraformaldehyde fixed after euthanasia and subsequently embedded in paraffin for hematoxylin-eosin (H&E) staining. Ovary sections of 5 µm thickness were deparaffinized initially for 20 minutes using xylene and processed and stained for hematoxylin-eosin (H&E) staining as detailed in(33) and imaged using Nikon microscope (CLIPSETi-S, Japan).

### Hormonal Analysis

Serum was obtained by centrifuging blood samples from both control and PMOS (DHEA-induced) groups at 1500xg for 10 minutes at 4°C. The samples were then stored at −80°C for subsequent hormonal analysis. ELISA kits from MyBioSource, San Diego, USA were employed for analyzing circulating levels of testosterone (MBS702281), luteinizing hormone (LH) (cat # MBS2514287), and follicle stimulating hormone (FSH) (MBS2507988). Procedures were conducted according to the manufacturer’s instructions.

### Statistical Analysis

The values for all variables were reported as mean ± SD. Depending on the data distribution, expression levels between the PMOS and control groups were compared using an unpaired Student’s t-test, a Mann–Whitney U test, or a Fisher-Pitman permutation test. Statistical analyses were performed using GraphPad Prism 9 and Excel’s Data Analysis ToolPak. P-values ≤ 0.05 were considered significant.

## Results

### DHEA treatment in mice resulted in PMOS morphology

Administration of DHEA in mice for 20 consecutive days resulted in the development of reproductive abnormalities. Initially, the control group had n=22 and DHEA group had n=21 mice. Testosterone levels were higher in the DHEA-treated group(n=15) compared to controls (P< 0.0001), after excluding 6 mice with DHEA levels > 2 SD below the mean of the control (Figure 2A). FSH levels did not show marked variation among the two groups (Figure 2B). LH levels were significantly increased among the DHEA-treated group compared to the sham-treated group (P<0.01) (Figure 2C), which is further reflected in the elevated LH/FSH ratio (P<0.05) (Figure2D). The ovaries of control mice displayed healthy morphology with growing follicles of all stages. DHEA-treated mice developed multiple cystic and atretic follicles in the ovaries (Figure 2E). Examination of the body weight of the DHEA-treated and placebo-treated (control) groups revealed no significant alteration in body weight at the end of the treatment (Figure 2F). Assessment of vaginal cytology displayed that the DHEA-treated mice were entirely acyclic and continued in the persistent estrus stage(Figure 2G). In contrast, mice from the control group had a regular oestrous cycle (Figure 2G). Therefore, DHEA treatment in mice developed hallmark PMOS reproductive traits such as cystic morphology of the ovary, irregular estrous cycle, and elevated serum testosterone and LH levels.

### DHEA-induced PMOS mice exhibited anxiety-like behaviour and impaired cognition

The experimental setup depicted in Figure 3A-B has been explained elsewhere. While there was no significant change in the time spent in the CZ(Figure 3C), DHEA-induced PMOS mice spent more time in corners of the arena (P<0.05) (Figure 3D), less time spent in the PZ (P<0.01)(Figure 3E) and, decreased number of entries to the CZ of the arena (P<0.01) (Figure 3F) when compared to the control group pointing to the anxiety-like behavior of mice with PMOS. There was no significant change in latency to the center(Figure 3G) and average speed (Figure 3H) among the groups. Treatment of mice with DHEA resulted in a decrease in total distance traveled inside the arena(P<0.05) (Figure 3I), indicating the depression-like behavior of DHEA-induced PMOS mice when compared to the control group.

### DHEA-treated PMOS mice showed deficits in reference memory

The experimental setup depicted in Figure 4A has been explained earlier. DHEA-treated mice also spent less time (P<0.05) with novel objects when compared to familiar objects based on the discrimination index test showing a defective Novel object recognition memory (Figure 4B). DHEA-treated mice displayed a significant decrease in object location memory with lesser time spent in the object at a novel location in comparison with an object in a familiar place (P<0.05) (Figure 4C), denoting the impairment of hippocampus-dependent spatial memory and recognition memory in DHEA-induced PMOS mice. There was no significant difference in the total time spent investigating the objects in either test (Fig. 4D & E), indicating that the observed deficits in object location memory were not due to alterations in exploratory drive.

### DHEA-induced PMOS mice show altered hippocampus expression of immediate early genes and long-term memory markers

DHEA-treated PMOS mice were evaluated for variation in hippocampal expression levels of immediate early genes like NPAS4, EGR1/2, and ARC and long-term memory markers like NMDAR1 and NMDAR2A. Transcript levels of *Npas4* were significantly reduced (P<0.05) in DHEA-induced PMOS mice with a fold change of (FC) = -2.3 when compared to controls (Figure 5A). This was further corroborated by our western blot data exhibiting reduced NPAS4 protein expression in PMOS (P<0.05) (Figure 5B-C) relative to their controls. Immunohistochemical localization revealed a distinct loss of NPAS4 expression in the hippocampus of PMOS model DHEA-treated mice when relative to the controls (P<0.0001) (Figure 5D-E). A stark reduction in hippocampal neurite processes is seen in cornu ammonis (CA1 and CA3) and dentate gyrus (DG) (Figure 5D).

**Figure 5:**
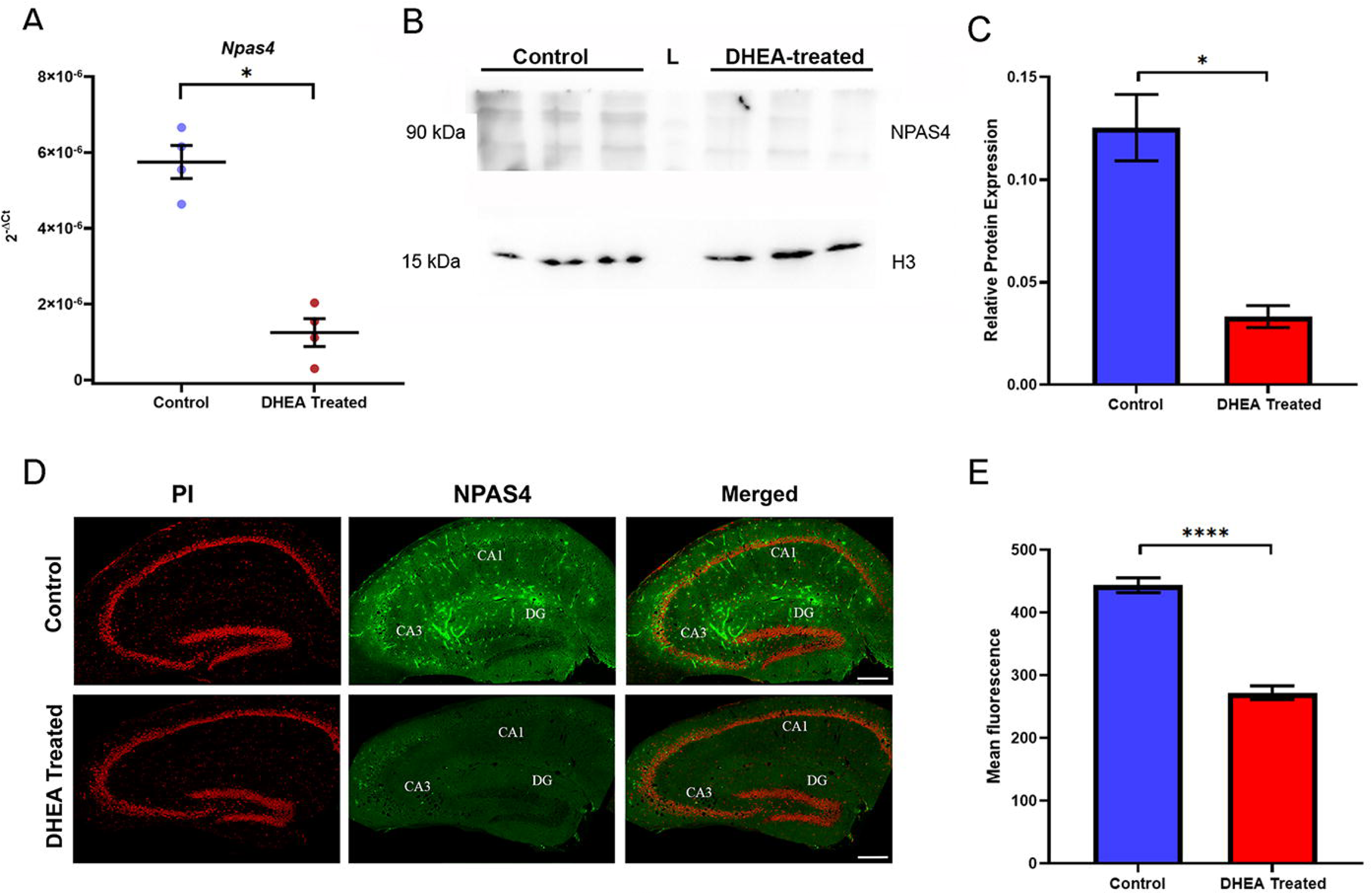
Expression of NPAS4 in hippocampus of DHEA-induced PMOS and control mice. A) Dot plot representation of relative mRNA level of *Npas4* in Control and DHEA treated mice hippocampus (n=4) as determined by q-RT PCR with 18SrRNA as the endogenous control B)Western blot analysis of NPAS4: in Control (lanes 1–3), DHEA-induced PMOS (lanes 5–7). Histone3 was used as the endogenous control for comparison. C) Densitometry analysis of NPAS4 is shown as a bar graph representation (right). D) Immunohistochemistry analysis showing NPAS4 expression in the hippocampus of Control and DHEA-treated mice. NPAS4 probed with the primary antibody was labelled with anti-mouse secondary DyLight 488, and PI was used as the nuclear stain. The sub-regions of the hippocampus, CA1 (Cornu Ammonis 1), CA3 (Cornu Ammonis 3), and DG (dentate gyrus), are marked. E)Region of interest (ROI) quantification of fluorescence intensity for NPAS4 in the hippocampus is shown as a bar graph representation. Control group (blue) and DHEA-treated group (brown) are shown as individual data points in the figure. Data presented as mean ±SEM. Mann-Whitney U was calculated for analysing statistical significance for graph A, and B, Fisher-Pitman permutation test for C, and Unpaired Student’s T-test for E. P-values <0.05 were set to be significant. *P-value <0.05, **P-value < 0.01, ***P-value <0.001 and ****P-value <0.0001

Expression levels of long-term memory marker *Nmdar1 (Grin1-(glutamate ionotropic receptor NMDA type subunit 1))* was significantly upregulated in DHEA-induced PMOS mice with a fold change of 4.7 when compared to controls (P<0.05) (Figure 6A). In contrast, *Nmdar2a (Grin2a)* transcript levels showed significant downregulation in DHEA-induced PMOS mice with a fold change of (−11.1) when compared to controls (P<0.05) (Figure 6B). Transcript levels of *Arc* showed a significant increase (P<0.05) between the control and experimental group (Figure 6C).

**Figure 6:**
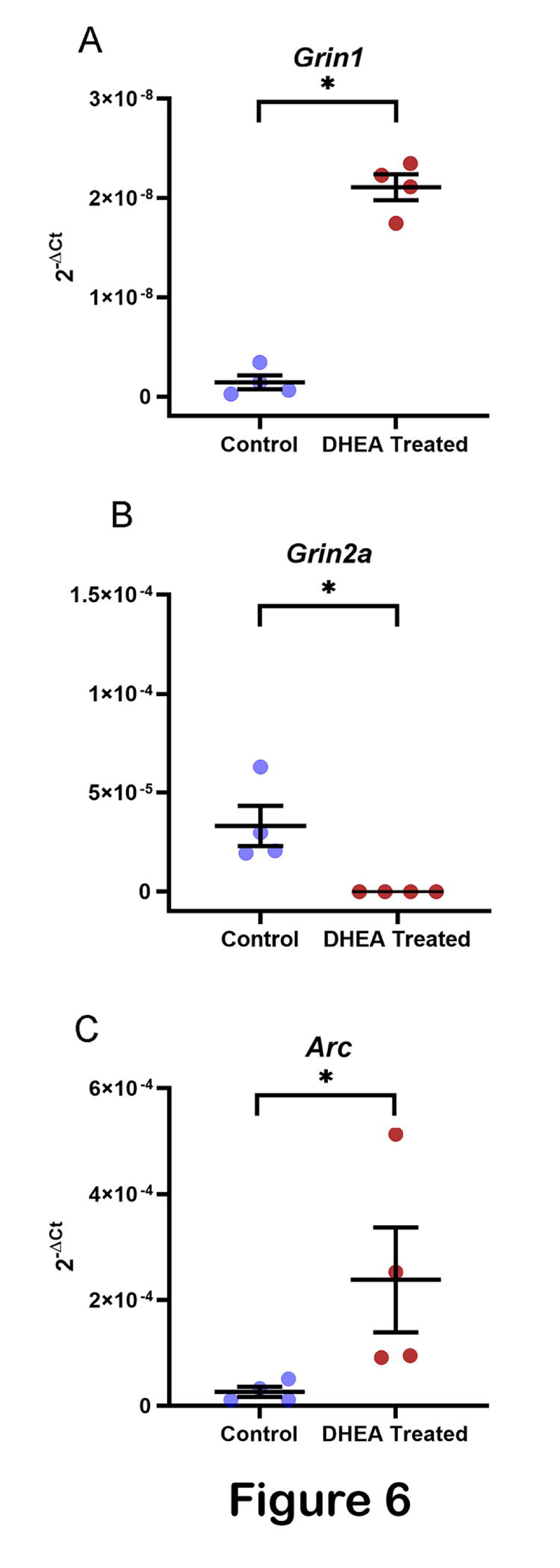
Expression analysis of Grin and Arc memory markers in the hippocampus of control and DHEA-induced PMOS mice. Dot plot representation of relative mRNA level of A) *Grin1,* B) *Grin2a,* and C) *Arc* in the hippocampus of control mice and DHEA-induced PMOS (n=4) as determined by q-RT PCR with 18S rRNA as the endogenous control. Control group (blue) and DHEA-treated group (brown) are shown as individual data points in the figure. Data presented as mean ±SEM. Mann-Whitney U was calculated for analysing statistical significance. P-values <0.05 were set to be significant. *P-value <0.05, **P-value < 0.01, ***P-value <0.001 and ****P-value <0.0001.

Egr1 and Egr2 expression levels were significantly higher in DHEA-induced PMOS mice, showing fold changes of 2.9 (P < 0.05) and 2.7 (P < 0.05), respectively, relative to the control group (Figure 7A-B). Significant upregulation of EGR1 protein levels was observed in the hippocampus of DHEA-treated mice relative to control mice (P < 0.05) (Figure 7C-D).

**Figure 7:**
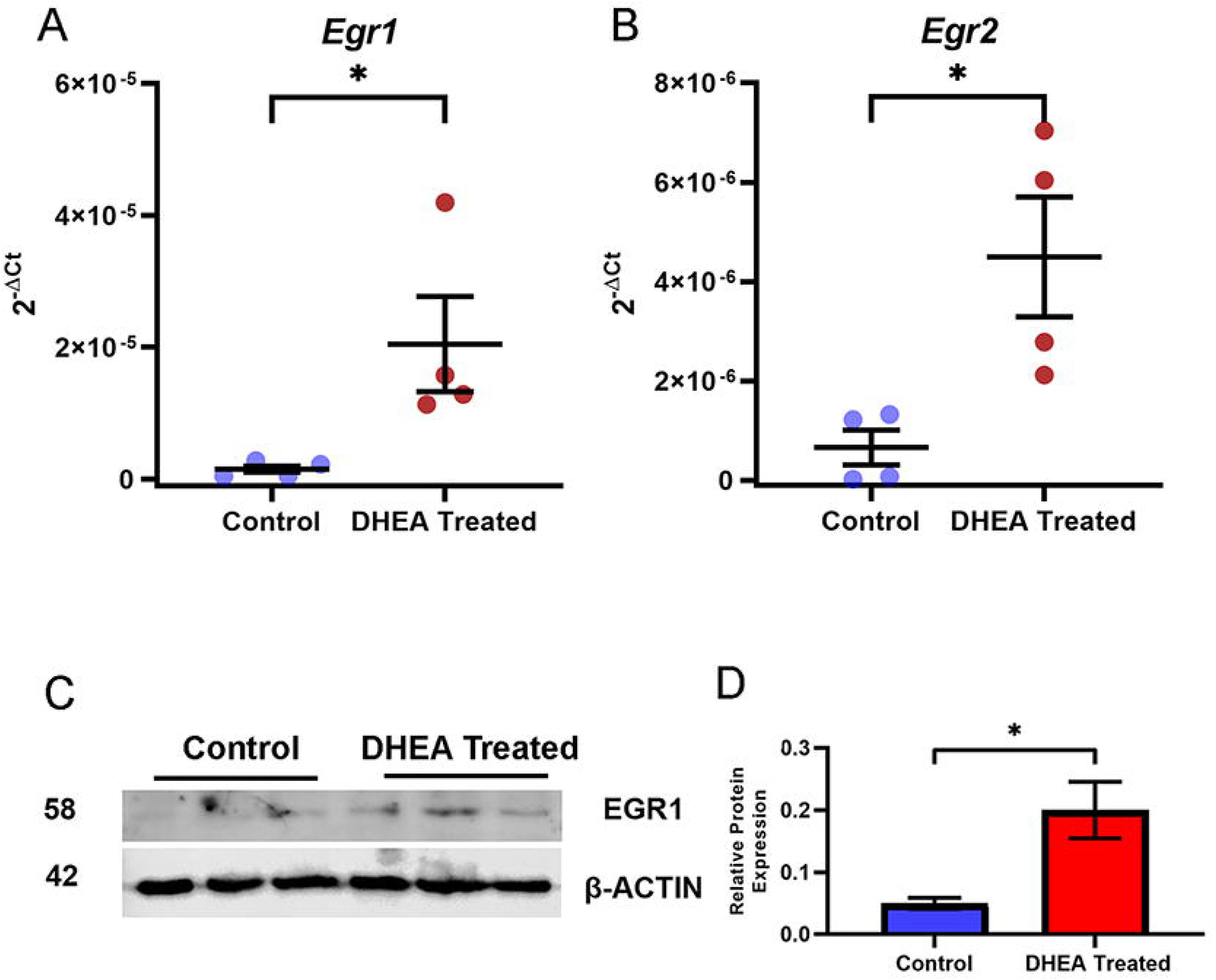
Expression analysis of *Egr1* and *Egr2* in the hippocampus of control and DHEA-induced PMOS mice. Dot plot representation of relative mRNA level of memory markers; A) *Egr1* and B) *Egr2* in the hippocampus of DHEA-induced PMOS and control mice (n=4) as determined by q-RT PCR with 18SrRNA as the endogenous control C) Western blot analysis of EGR1 in Control (lanes 1–3), DHEA-induced PMOS (lanes 4–6). β-actin was used as the endogenous control for comparison. D) Densitometry analysis of EGR1 is shown as a bar graph representation. Control group (blue) and DHEA-treated group (brown) are shown as individual data points in the figure. Data presented as mean ±SEM. Mann-Whitney U was calculated for analysing statistical significance for graph A and B and a Fisher-Pitman permutation test for D. P-values <0.05 were set to be significant. *P-value <0.05, **P-value < 0.01, ***P-value <0.001 and ****P-value <0.0001.

### DHEA-induced PMOS mice show altered hippocampus expression of Androgen receptor (AR)

The exact etiology of PMOS remains unclear, but recent studies suggest a link to the androgen receptor (AR) in the brain to the disease. We investigated how AR levels in the hippocampus, a region critical for cognition and memory, are altered in PMOS-like conditions. Expression levels of *Androgen Receptor (Ar)* were significantly upregulated in the hippocampus of DHEA-induced PMOS mice with a fold change of 1.9 when compared to controls (P<0.05) (Figure 8).

**Figure 8:**
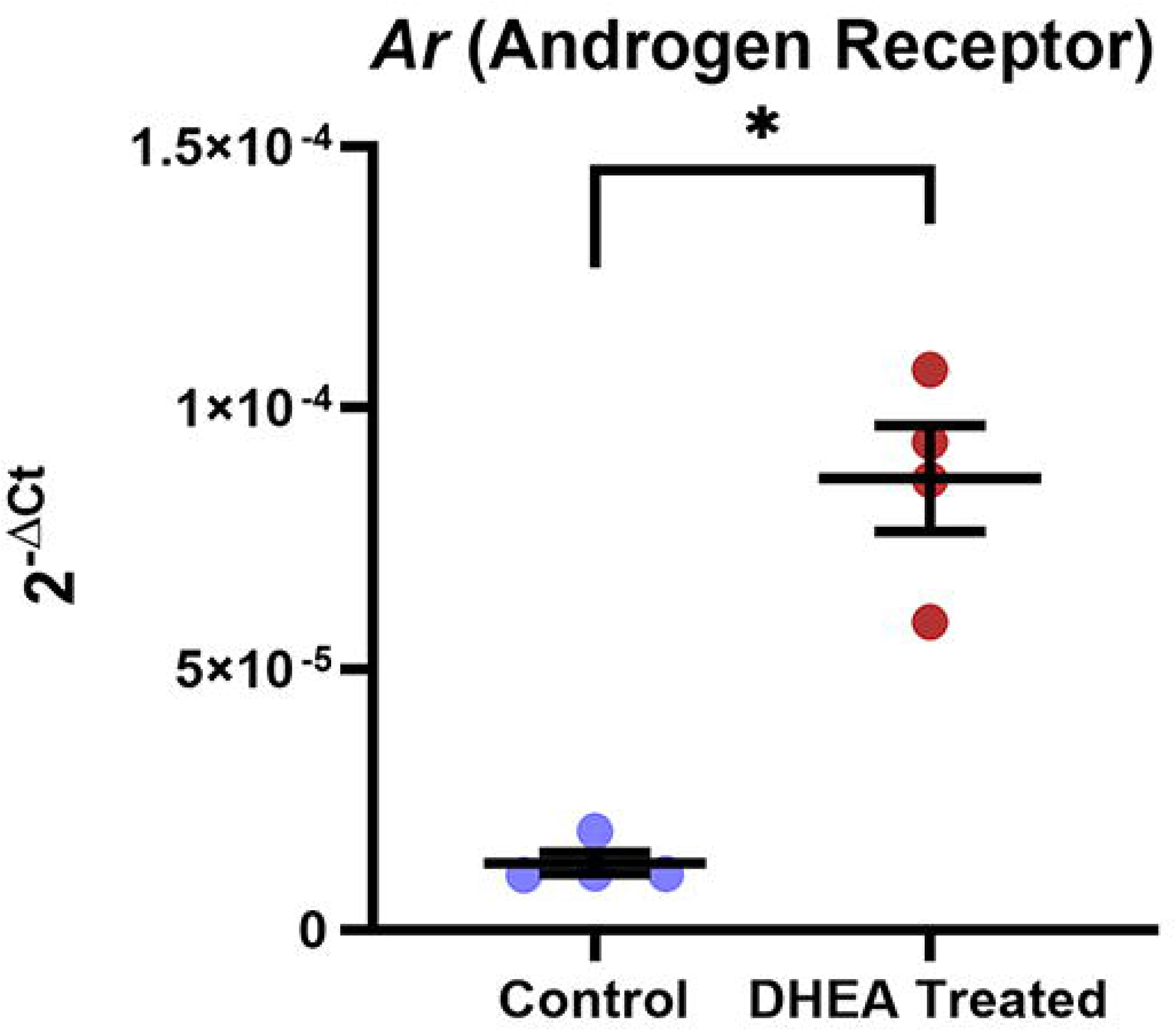
Androgen Receptor expression analysis in the hippocampus of control and DHEA-induced PMOS mice. Dot plot representation of relative mRNA level of *AR (Androgen receptor)* in the hippocampus of control mice and DHEA-induced PMOS (n=4) as determined by q-RT PCR with 18SrRNA as the endogenous control. Data presented as mean ±SEM. Control group (blue) and DHEA-treated group (brown) are shown as individual data points in the figure. Mann-Whitney U was calculated for analysing statistical significance. P-values <0.05 were set to be significant. *P-value <0.05.

## Discussion

Sex steroids and metabolic factors play a key role in shaping sex-specific differences in cognitive ability(40). Since PMOS is a disorder that affects both the reproductive and metabolic systems(41), it presents an unique opportunity to study the effects of a multisystemic burden on cognition proficiency.

Major depressive disorder and generalized anxiety disorder are associated with higher DHEAS levels in PMOS women(42). This burden is further exacerbated by lifestyle disruptions common in PMOS, including sleep disorders(43), particularly Obstructive Sleep Apnea(OSA)(44) which is strongly correlated with anxiety and depression in PMOS(45). This reinforces the significant prevalence of psychological conditions, including depression and anxiety disorders recorded among women with PMOS(46; 47). Anxiety-like behavior is reported in DHT-induced PMOS in female rats as well(48). Our open-field test(OFT) data shows that DHEA-treated mice spent less time in the central zone and total distance traveled (Figure 3), indicating depression and anxiety-like behavior in the androgen-induced PMOS mice model. Our results corroborate earlier findings and suggest that androgen-induced animal models mimic mood disturbances seen in women with PMOS.

Interestingly, it has been proposed that hippocampus may have a role in anxiety and memory(49). The hippocampus of the brain is known to store information about places and their spatial organizations in an organism’s environment, as well as the presence of objects at particular locations. Therefore, the hippocampus is part of one’s memory unit and supports a learning map of the outside world(50). Our examination of whether hippocampal dependent-memory defects are also displayed along with depression and anxiety-like behavior in DHEA-treated mice by employing OLT and NORT reveals that DHEA-induced PMOS mice spent significantly less time exploring the object in novel spatial location compared to the familiar one (Figure 4C). DHEA-treated mice showed reduced preference for a novel object over a familiar object when compared to the control group implicating a memory dysfunction associated with long-term object recognition under hyperandrogenic state (Figure 4B). As the NORT depends on interactions between the hippocampus and perirhinal cortex which is involved in processing visual stimuli and transmitting them to hippocampus; our findings further support disrupted memory processing in DHEA-induced PMOS mice. Our results align with the findings of Bussey et al who reported that object recognition deficits can arise from dysfunction in hippocampal input–output pathways such as the fimbria–fornix(51). Neuroimaging of the brain using resting state-fMRI in women with PMOS showed that abnormal brain function at specific regions and network levels with an association with serum hormones and cognition(52). Thus, our findings highlight the need of in-depth brain imaging of PMOS women particularly focusing on hippocampus, perirhinal cortex, and associated white-matter to better understand the basis of recognition memory dysfunction.

*Npas4*, an immediate early gene, directs activity-dependent gene expression in neurons to promote circuit activity, and *Npas4* knockouts have altered neuronal plasticity(53). Our observation of significantly reduced hippocampal *Npas4* transcript and NPAS4 protein expression points to the role of NPAS4 in hippocampal object memory dysfunction in DHEA-induced PMOS mice (Figure 5). Our finding of reduced NPAS4-positive neurites in PMOS hippocampus is consistent with the evidence that decreased NPAS4 in the dentate gyrus(DG) impairs neurogenesis and hippocampal-dependent memory(54). Increased neurogenesis in the hippocampus is reported to be essential for navigational learning and memory storage(55).

Interestingly, *Grin1,* the obligatory GluN1 NMDA receptor subunit, was elevated (Figure 6A) which codes for the obligatory GluN1 NMDA receptor subunit suggests an overexpression of *Grin1* under an androgenic stimulus in the DHEA-treated PMOS mouse model. Glutamate is one of the ligands for NMDA receptor and elevated glutamate levels in the hippocampus of mouse models of PMOS are reported(56) highlighting its link to brain alterations in this condition. It is also to be remembered that glutamatergic input is known to be important for reward-seeking learning. Along with this, *Arc*, a bidirectional plasticity regulator, is also upregulated in our study, as seen before in high-glutamatergic environments(57). In contrast, a reduced *Grin2a* expression in DHEA-induced PMOS mouse (Figure 6B) could lead to improper NMDAR receptor assembly and may be a critical determinant of the observed spatial memory deficits, as *Grin2a* knockdown is linked to impairments in spatial memory and cognition(58). Together with glutamatergic excess, these alterations would synergize to impair hippocampal plasticity and memory under hyperandrogenic conditions.

*Egr1* is another key regulator of *Grin1*, since chromatin immunoprecipitation studies in brain have identified *Grin1* as a bonafide target gene of EGR1(*59*). Our results reveal an overexpression of memory markers *Egr1 and Egr2* in the hippocampus of DHEA-treated PMOS mice (Figure 7). Thus, the increased EGR1 could modulate *Grin1* expression as observed in our studies and this is in line with reports of Koldamova et al(59). EGR1 is essential for long-term memory consolidation in the hippocampus, but remarkably, aberrant increases of EGR1 activity observed in superficial CA1 pyramidal neurons under chronic stress are associated with marked spatial learning deficits(60). Reduction of EGR1-positive CA1 neuronal ensembles is associated with improved motor learning while upregulation of Egr1 causes increased CA1 neuronal assembles which limit motor learning ability on rotarod task(61). Additionally, *Egr2* mutant mice in addition to improved object recognition memory, have superior motor learning ability(60). Thus, increased combinatorial impact of elevated *Egr1* and *Egr2* in our study suggest a synergistic impact on motor learning ability in PMOS. Given the limited understanding of motor learning in PMOS, our findings highlight an important area for future investigation.

Androgen receptors in the CA1 of the hippocampus are important for visual information processing via NMDAR activity in mice(62). Evidence suggests that AR activity must remain within an optimal range to support NMDAR-dependent LTP, as both excessive and insufficient signalling impair this process(63). In PMOS, chronically elevated androgen levels may push AR signalling outside this physiological window, altering NMDAR function and impairing CA1 plasticity. In line with this, our observation of increased AR expression in the hippocampus (Figure 8) suggests that dysregulated AR signalling may contribute to the cognitive difficulties reported in PMOS.

Our study is the first to report differential memory and learning ability in mice models of PMOS orchestrated by decreased NPAS4-Grin2a axis alonside increased *Grin1, Arc* and *Egr1/2*. We acknowledge that collecting samples at a single time point is a limitation and does not capture peak dynamics, and future work should assess the temporal dynamics of immediate early gene expression to provide a more complete understanding of memory and learning in PMOS. In summary, our findings demonstrate that PMOS-like conditions disrupt the brain’s ability to support memory formation, as evidenced by behavioural tests and altered markers of neural activity. This highlights the critical impact of hormonal dysregulation on cognitive function in PMOS and underscores the need for targeted interventions to mitigate its neurological consequences. Future clinical studies for validating these observations in PMOS women are recommended as PMOS-related cognitive dysfunction is an emerging field of study with far-reaching future public health implications.

## Data availability

The data generated for the study underlying this article are available in the article and in its online supplementary material. The data will be shared on reasonable request to the corresponding author.

## Declaration of Interest

The authors declare that there is no conflict of interest that could be perceived as prejudicing the impartiality of the research reported.

## Funding

This work was supported by the RGCB-DBT Core Funds to M.L. M. L. is an ICMR Emeritus Scientist (ICMR Letter No. HRD/Head/IES/2024/9). B.S.J. was supported by a Research Fellowship from Indian Council of Medical Research (RBMH/FW/2020/2). S. R. was supported by a fellowship under the SSR Scheme of DST-SERB funds (Grant no. CRG/2019/005029) to M.L.

## Author contribution statement

M.L. was involved in the project conception and supervision, experiment design, and manuscript review and editing. S.R. designed Fig. 1 B.S.J. and S.R. designed and executed the experiments in Figs 2–7, analyzed data, and wrote the manuscript draft. The graphical summary was designed by M.L. and S.R. All authors discussed the results, commented on the manuscript and approved the final version.

## Acknowledgments

The authors acknowledge the support extended by Dr. R. V. Omkumar for allowing the use of facility for behavioral experiments in the Neurobiology Division of RGCB.. The support by RGCB Animal Research Facility staff during animal experiments particularly the help with perfusion of mice provided by Dr. Vishnu Sunil Jayakumar is acknowledged.

## Notes

### Competing Interest Statement

The authors have declared no competing interest.

### Summary of Updates

Following a landmark global consensus, the condition previously known as Polycystic Ovary Syndrome (PCOS) has been officially renamed Polyendocrine Metabolic Ovarian Syndrome (PMOS) and hence in this revised version, we have updated this nomenclature in the Manuscript file, Figure 1, and Graphical Abstract.

## References

1. Franks S 1995 Polycystic ovary syndrome. N Engl J Med 333 853–861. (10.1056/NEJM199509283331307)

2. Stein IF 1955 [The Stein-Leventhal syndrome]. Rev Obstet Ginecol Venez 15 699–713.

3. Rotterdam EA-SPcwg 2004 Revised 2003 consensus on diagnostic criteria and long-term health risks related to polycystic ovary syndrome (PMOS). Hum Reprod 19 41–47. (10.1093/humrep/deh098)

4. Wolf WM, Wattick RA, Kinkade ON, et al. 2018 Geographical Prevalence of Polycystic Ovary Syndrome as Determined by Region and Race/Ethnicity. Int J Environ Res Public Health 15. (10.3390/ijerph15112589)

5. Chang RJ, Nakamura RM, Judd HL, et al. 1983 Insulin resistance in nonobese patients with polycystic ovarian disease. J Clin Endocrinol Metab 57 356–359. (10.1210/jcem-57-2-356)

6. Dokras A 2013 Cardiovascular disease risk in women with PMOS. Steroids 78 773–776. (10.1016/j.steroids.2013.04.009)

7. Eggers S & Kirchengast S 2001 The polycystic ovary syndrome--a medical condition but also an important psychosocial problem. Coll Antropol 25 673–685.

8. Elsenbruch S, Hahn S, Kowalsky D, et al. 2003 Quality of life, psychosocial well-being, and sexual satisfaction in women with polycystic ovary syndrome. J Clin Endocrinol Metab 88 5801–5807. (10.1210/jc.2003-030562)

9. Moran LJ, Deeks AA, Gibson-Helm ME, et al. 2012 Psychological parameters in the reproductive phenotypes of polycystic ovary syndrome. Hum Reprod 27 2082–2088. (10.1093/humrep/des114)

10. Mansson M, Holte J, Landin-Wilhelmsen K, et al. 2008 Women with polycystic ovary syndrome are often depressed or anxious--a case control study. Psychoneuroendocrinology 33 1132–1138. (10.1016/j.psyneuen.2008.06.003)

11. Rassi A, Veras AB, dos Reis M, et al. 2010 Prevalence of psychiatric disorders in patients with polycystic ovary syndrome. Compr Psychiatry 51 599–602. (10.1016/j.comppsych.2010.02.009)

12. Cesta CE, Mansson M, Palm C, et al. 2016 Polycystic ovary syndrome and psychiatric disorders: Co-morbidity and heritability in a nationwide Swedish cohort. Psychoneuroendocrinology 73 196–203. (10.1016/j.psyneuen.2016.08.005)

13. Rees DA, Udiawar M, Berlot R, et al. 2016 White Matter Microstructure and Cognitive Function in Young Women With Polycystic Ovary Syndrome. J Clin Endocrinol Metab 101 314–323. (10.1210/jc.2015-2318)

14. Huddleston HG, Jaswa EG, Casaletto KB, et al. 2024 Associations of Polycystic Ovary Syndrome With Indicators of Brain Health at Midlife in the CARDIA Cohort. Neurology 102 e208104. (10.1212/WNL.0000000000208104)

15. Ruddenklau A & Campbell RE 2019 Neuroendocrine Impairments of Polycystic Ovary Syndrome. Endocrinology 160 2230–2242. (10.1210/en.2019-00428)

16. Coutinho EA & Kauffman AS 2019 The Role of the Brain in the Pathogenesis and Physiology of Polycystic Ovary Syndrome (PMOS). Med Sci (Basel) 7. (10.3390/medsci7080084)

17. Hyde Z, Flicker L, Almeida OP, et al. 2010 Higher luteinizing hormone is associated with poor memory recall: the health in men study. J Alzheimers Dis 19 943–951. (10.3233/JAD-2010-1342)

18. Lai W, Li X, Zhu H, et al. 2020 Plasma luteinizing hormone level affects the brain activity of patients with polycystic ovary syndrome. Psychoneuroendocrinology 112 104535. (10.1016/j.psyneuen.2019.104535)

19. Luine VN 2008 Sex steroids and cognitive function. J Neuroendocrinol 20 866–872. (10.1111/j.1365-2826.2008.01710.x)

20. Sanders G, Sjodin M & de Chastelaine M 2002 On the elusive nature of sex differences in cognition: hormonal influences contributing to within-sex variation. Arch Sex Behav 31 145–152. (10.1023/a:1014095521499)

21. Hogervorst E, Matthews FE & Brayne C 2010 Are optimal levels of testosterone associated with better cognitive function in healthy older women and men? Biochim Biophys Acta 1800 1145–1152. (10.1016/j.bbagen.2009.12.009)

22. Soleman RS, Kreukels BPC, Veltman DJ, et al. 2016 Does polycystic ovary syndrome affect cognition? A functional magnetic resonance imaging study exploring working memory. Fertil Steril 105 1314–1321 e1311. (10.1016/j.fertnstert.2016.01.034)

23. Schattmann L & Sherwin BB 2007 Testosterone levels and cognitive functioning in women with polycystic ovary syndrome and in healthy young women. Horm Behav 51 587–596. (10.1016/j.yhbeh.2007.02.007)

24. Barry JA, Parekh HS & Hardiman PJ 2013 Visual-spatial cognition in women with polycystic ovarian syndrome: the role of androgens. Hum Reprod 28 2832–2837. (10.1093/humrep/det335)

25. Wei W, Chen C, Dong Q, et al. 2016 Sex Differences in Gray Matter Volume of the Right Anterior Hippocampus Explain Sex Differences in Three-Dimensional Mental Rotation. Front Hum Neurosci 10 580. (10.3389/fnhum.2016.00580)

26. Epstein RA, Patai EZ, Julian JB, et al. 2017 The cognitive map in humans: spatial navigation and beyond. Nat Neurosci 20 1504–1513. (10.1038/nn.4656)

27. Lee SLT, Lew D, Wickenheisser V, et al. 2019 Interdependence between dorsal and ventral hippocampus during spatial navigation. Brain Behav 9 e01410. (10.1002/brb3.1410)

28. Tabori NE, Stewart LS, Znamensky V, et al. 2005 Ultrastructural evidence that androgen receptors are located at extranuclear sites in the rat hippocampal formation. Neuroscience 130 151–163. (10.1016/j.neuroscience.2004.08.048)

29. Caldwell ASL, Edwards MC, Desai R, et al. 2017 Neuroendocrine androgen action is a key extraovarian mediator in the development of polycystic ovary syndrome. Proc Natl Acad Sci U S A 114 E3334–E3343. (10.1073/pnas.1616467114)

30. Yu Q, Hao S, Wang H, et al. 2016 Depression-Like Behavior in a Dehydroepiandrosterone-Induced Mouse Model of Polycystic Ovary Syndrome. Biol Reprod 95 79. (10.1095/biolreprod.116.142117)

31. Antunes M & Biala G 2012 The novel object recognition memory: neurobiology, test procedure, and its modifications. Cogn Process 13 93–110. (10.1007/s10339-011-0430-z)

32. Ennaceur A 2010 One-trial object recognition in rats and mice: methodological and theoretical issues. Behav Brain Res 215 244–254. (10.1016/j.bbr.2009.12.036)

33. Johnson BS, Krishna MB, Padmanabhan RA, et al. 2022 Derailed peripheral circadian genes in polycystic ovary syndrome patients alters peripheral conversion of androgens synthesis. Hum Reprod 37 1835–1855. (10.1093/humrep/deac139)

34. Vandresen-Filho S, Franca LM, Alcantara-Junior J, et al. 2015 Statins enhance cognitive performance in object location test in albino Swiss mice: involvement of beta-adrenoceptors. Physiol Behav 143 27–34 (10.1016/j.physbeh.2015.02.024)

35. Lueptow LM 2017 Novel Object Recognition Test for the Investigation of Learning and Memory in Mice. J Vis Exp. (10.3791/55718)

36. Sik A, van Nieuwehuyzen P, Prickaerts J, et al. 2003 Performance of different mouse strains in an object recognition task. Behav Brain Res 147 49–54. (10.1016/s0166-4328(03)00117-7)

37. Denninger JK, Smith BM & Kirby ED 2018 Novel Object Recognition and Object Location Behavioral Testing in Mice on a Budget. J Vis Exp. (10.3791/58593)

38. Hattiangady B, Mishra V, Kodali M, et al. 2014 Object location and object recognition memory impairments, motivation deficits and depression in a model of Gulf War illness. Front Behav Neurosci 8 78. (10.3389/fnbeh.2014.00078)

39. Rueden CT, Schindelin J, Hiner MC, et al. 2017 ImageJ2: ImageJ for the next generation of scientific image data. BMC Bioinformatics 18 529. (10.1186/s12859-017-1934-z)

40. Gurvich C, Hoy K, Thomas N, et al. 2018 Sex Differences and the Influence of Sex Hormones on Cognition through Adulthood and the Aging Process. Brain Sci 8. (10.3390/brainsci8090163)

41. Teede H, Deeks A & Moran L 2010 Polycystic ovary syndrome: a complex condition with psychological, reproductive and metabolic manifestations that impacts on health across the lifespan. BMC Med 8 41. (10.1186/1741-7015-8-41)

42. Annagur BB, Tazegul A, Uguz F, et al. 2013 Biological correlates of major depression and generalized anxiety disorder in women with polycystic ovary syndrome. J Psychosom Res 74 244–247. (10.1016/j.jpsychores.2013.01.002)

43. Moran LJ, March WA, Whitrow MJ, et al. 2015 Sleep disturbances in a community-based sample of women with polycystic ovary syndrome. Hum Reprod 30 466–472. (10.1093/humrep/deu318)

44. Fogel RB, Malhotra A, Pillar G, et al. 2001 Increased prevalence of obstructive sleep apnea syndrome in obese women with polycystic ovary syndrome. J Clin Endocrinol Metab 86 1175–1180. (10.1210/jcem.86.3.7316)

45. Zhou X, Jaswa E, Pasch L, et al. 2021 Association of obstructive sleep apnea risk with depression and anxiety symptoms in women with polycystic ovary syndrome. J Clin Sleep Med 17 2041–2047. (10.5664/jcsm.9372)

46. Dybciak P, Humeniuk E, Raczkiewicz D, et al. 2022 Anxiety and Depression in Women with Polycystic Ovary Syndrome. Medicina (Kaunas) 58. (10.3390/medicina58070942)

47. Jedel E, Waern M, Gustafson D, et al. 2010 Anxiety and depression symptoms in women with polycystic ovary syndrome compared with controls matched for body mass index. Hum Reprod 25 450–456. (10.1093/humrep/dep384)

48. Feng Y, Shao R, Weijdegard B, et al. 2011 Effects of androgen and leptin on behavioral and cellular responses in female rats. Horm Behav 60 427–438. (10.1016/j.yhbeh.2011.07.012)

49. Bannerman DM, Sprengel R, Sanderson DJ, et al. 2014 Hippocampal synaptic plasticity, spatial memory and anxiety. Nat Rev Neurosci 15 181–192. (10.1038/nrn3677)

50. Manns JR & Eichenbaum H 2009 A cognitive map for object memory in the hippocampus. Learn Mem 16 616–624. (10.1101/lm.1484509)

51. Bussey TJ, Duck J, Muir JL, et al. 2000 Distinct patterns of behavioural impairments resulting from fornix transection or neurotoxic lesions of the perirhinal and postrhinal cortices in the rat. Behav Brain Res 111 187–202. (10.1016/s0166-4328(00)00155-8)

52. Li G, Hu J, Zhang S, et al. 2020 Changes in Resting-State Cerebral Activity in Women With Polycystic Ovary Syndrome: A Functional MR Imaging Study. Front Endocrinol (Lausanne) 11 603279. (10.3389/fendo.2020.603279)

53. Bloodgood BL, Sharma N, Browne HA, et al. 2013 The activity-dependent transcription factor NPAS4 regulates domain-specific inhibition. Nature 503 121–125. (10.1038/nature12743)

54. Yun J, Koike H, Ibi D, et al. 2010 Chronic restraint stress impairs neurogenesis and hippocampus-dependent fear memory in mice: possible involvement of a brain-specific transcription factor Npas4. J Neurochem 114 1840–1851. (10.1111/j.1471-4159.2010.06893.x)

55. Berdugo-Vega G, Arias-Gil G, Lopez-Fernandez A, et al. 2020 Increasing neurogenesis refines hippocampal activity rejuvenating navigational learning strategies and contextual memory throughout life. Nat Commun 11 135. (10.1038/s41467-019-14026-z)

56. Chaudhari N, Dawalbhakta M & Nampoothiri L 2018 GnRH dysregulation in polycystic ovarian syndrome (PMOS) is a manifestation of an altered neurotransmitter profile. Reprod Biol Endocrinol 16 37. (10.1186/s12958-018-0354-x)

57. Waung MW, Pfeiffer BE, Nosyreva ED, et al. 2008 Rapid translation of Arc/Arg3.1 selectively mediates mGluR-dependent LTD through persistent increases in AMPAR endocytosis rate. Neuron 59 84–97. (10.1016/j.neuron.2008.05.014)

58. Du Z, Song Y, Chen X, et al. 2021 Knockdown of astrocytic Grin2a aggravates beta-amyloid-induced memory and cognitive deficits through regulating nerve growth factor. Aging Cell 20 e13437. (10.1111/acel.13437)

59. Koldamova R, Schug J, Lefterova M, et al. 2014 Genome-wide approaches reveal EGR1-controlled regulatory networks associated with neurodegeneration. Neurobiol Dis 63 107–114. (10.1016/j.nbd.2013.11.005)

60. Poirier R, Cheval H, Mailhes C, et al. 2007 Paradoxical role of an Egr transcription factor family member, Egr2/Krox20, in learning and memory. Front Behav Neurosci 1 6. (10.3389/neuro.08.006.2007)

61. Brito V, Montalban E, Sancho-Balsells A, et al. 2022 Hippocampal Egr1-Dependent Neuronal Ensembles Negatively Regulate Motor Learning. J Neurosci 42 5346–5360. (10.1523/JNEUROSCI.2258-21.2022)

62. Pouliot WA, Handa RJ & Beck SG 1996 Androgen modulates N-methyl-D-aspartate-mediated depolarization in CA1 hippocampal pyramidal cells. Synapse 23 10–19. (10.1002/(SICI)1098-2396(199605)23:1<10::AID-SYN2>3.0.CO;2-K)

63. Picot M, Billard JM, Dombret C, et al. 2016 Neural Androgen Receptor Deletion Impairs the Temporal Processing of Objects and Hippocampal CA1-Dependent Mechanisms. PLoS One 11 e0148328. (10.1371/journal.pone.0148328)

