## Supplemental Table 1 for "Defective Hippocampus-Dependent Spatial Memory in Mouse Model of Polyendocrine metabolic ovarian syndrome (PMOS, formerly PCOS)"

Supplementary Table 1

List of primers for Real Time PCR:

| Sl. No. | Gene Symbol | Sequence | Length |
| --- | --- | --- | --- |
| 1 | m *Npas4* F | GATGCTGATCGCCTTTTCCG | 20 |
| 2 | m *Npas4* R | CAGGGTTTCCTGCCCAGTAG | 20 |
| 3 | m *Egr1* F | CCTGACCACAGAGTCCTTTTC | 21 |
| 4 | m *Egr1* R | GAGAAGCGGCCAGTATAGGTG | 21 |
| 5 | m *Egr2* F | GCGGGAGATGGCATGATCAAC | 21 |
| 6 | m *Egr2* R | ACCAGGGTACTGTGGGTCAA | 20 |
| 7 | m *Arc* F | TACCGTTAGCCCCTATGCCATC | 22 |
| 8 | m *Arc* R | TGATATTGCTGAGCCTCAACTG | 22 |
| 9 | m *Grin1* F | TCAGTGTGTGAGGACCTCATCTCT | 24 |
| 10 | m *Grin1* R | GAGTGAAGTGGTCGTTGGGAGTA | 23 |
| 11 | m *Grin2A* F | TAGACCTTAGCAGGCCCTCTC | 21 |
| 12 | m *Grin2A* R | GAGCTTTTGTTCCCCAAGAGT | 21 |
| 13 | m *Ar* F | CCTTGGATGGAGAACTACTCCG | 22 |
| 14 | m *Ar* R | TCCGTAGTGACAGCCAGAAGCT | 22 |
